# Preclinical assessment of MAGMAS inhibitor as a potential therapy for pediatric medulloblastoma

**DOI:** 10.1101/2024.02.29.582709

**Authors:** Zahra Motahari, Javier J. Lepe, Malia R. Bautista, Clay Hoerig, Ashley S. Plant-Fox, Bhaskar Das, Christie D. Fowler, Suresh N. Magge, Daniela A. Bota

## Abstract

Medulloblastoma, the most common pediatric brain malignancy, has Sonic Hedgehog (SHH) and non-SHH group3 subtypes. MAGMAS (Mitochondrial Associated Granulocyte Macrophage colony-stimulating factor Signaling molecules) encode for mitochondrial import inner membrane translocase subunit and is responsible for translocation of matrix proteins across the inner membrane. We previously reported that a small molecule MAGMAS inhibitor, BT9, decreases cell proliferation, migration, and oxidative phosphorylation in adult glioblastoma cell lines. The aim of our study was to investigate whether the chemotherapeutic effect of BT9 can be extended to pediatric medulloblastoma.

**Methods:** Multiple in vitro assays were performed using human DAOY (SHH activated tp53 mutant) and D425 (non-SHH group 3) cells. The impact of BT9 on cellular growth, death, migration, invasion, and metabolic activity were quantified using MTT assay, TUNEL staining, scratch wound assay, Matrigel invasion chambers, and seahorse assay, respectively. Survival following 50mg/kg BT9 treatment was assessed *in vivo* in immunodeficient mice intracranially implanted with D425 cells.

**Results:** Compared to control, BT9 treatment led to a significant reduction in medulloblastoma cell growth (DAOY, 24hrs IC50: 3.6uM, 48hrs IC50: 2.3uM, 72hrs IC50: 2.1uM; D425 24hrs IC50: 3.4uM, 48hrs IC50: 2.2uM, 72hrs IC50: 2.1uM) and a significant increase in cell death (DAOY, 24hrs p=0.0004, 48hrs p<0.0001; D425, 24hrs p=0.0001, 48hrs p=0.02). In DAOY cells, 3uM BT9 delayed migration, and significantly decreased DAOY and D425 cells invasion (p < 0.0001). Our *in vivo* study, however, did not extend survival in xenograft mouse model of group3 medulloblastoma compared to vehicle-treated controls.

**Conclusions:** Our *in vitro* data showed BT9 antitumor efficacy in DAOY and D425 cell lines suggesting that BT9 may represent a promising targeted therapeutic in pediatric medulloblastoma. These data, however, need to be further validated in animal models.

## Introduction

Medulloblastoma (MB) represents the most common pediatric brain malignancy and occurs mainly in children aged between 3 and 9 years [1, 2]. Standard therapy for MB includes surgical resection, craniospinal irradiation, and a combination of chemotherapeutic agents [3]. These treatments, however, can lead to severe side effects such as growth failure and cognitive deficits, and subsequent untreatable secondary malignancies [3]. Drug resistance is also very common, which severely limits the effectiveness of current chemotherapy [1]. Therefore, a more effective treatment approach is required to overcome the development of drug resistance and to improve clinical outcomes.

Medulloblastoma is a heterogenous disease and is categorized into WNT-activated, SHH-activated and *TP53*-mutant, SHH-activated and *TP53*-wild-type, and non-WNT/non-SHH groups. The latter group can be further subdivided to provisional subclasses, group 3 (G3) and group 4 (G4) when the ability to distinguish the two is possible [4]. Although each group has a different molecular signature, aberrant MYC proto-oncogenes (C-, N-, and L-) amplification seems a common theme in SHH, G3, and G4 [5]. N-MYC is highly expressed in SHH, G3, and G4, and C-MYC shows overexpression in the most aggressive G3 and G4 [5]. MYC proteins are considered the master regulators of cellular programs such as proliferation, apoptosis, and cell size. MYCs control metabolism through their regulatory effects of glycolytic genes such as hexokinase and lactate dehydrogenase [6]. In addition, MYC have been reported to regulate mitochondrial biogenesis by inducing genes involved in inner membrane, carboxylic acid metabolism, oxidative phosphorylation, and mtDNA replication [7-9]. The inner membrane proteins upregulated by MYC include TIM8 (translocase of the inner mitochondrial membrane), TIM10, TIM13, TIM17A, TIM22, TIM23, TIM44, and TIM16 or MAGMAS [7, 10]. As MYC is one of the most “undruggable” targets in cancer biology, alternative approaches targeting downstream pathways might prove useful on the -dependent malignancies, including MB [11].

TIM16 or MAGMAS (Mitochondrial Associated Granulocyte Macrophage colony-stimulating factor Signaling molecules), one of the MYC-regulated protein, is an ortholog of yeast pam16 (<u>p</u>resequence translocase-<u>a</u>ssociated protein import <u>m</u>otor) and is highly conserved in eukaryotes [9]. It encodes for mitochondrial import inner membrane translocase subunit TIM16. MAGMAS along with its partners (TIM17, TIM18, TIM44, Hsp70, Mge1) is an integral component of TIM23 import complex (TIM17, TIM23, TIM50) that together are responsible for translocation of matrix proteins across the inner membrane in an ATP-dependent process [12, 13]. MAGMAS controls the production of Reactive Oxygen Species (ROS) in the cells [14]. Overexpression of MAGMAS leads to reduction in ROS and increased cellular tolerance to oxidative stress, whereas its downregulation increases cellular ROS level and makes the cells more susceptible to ROS-mediated apoptosis [14, 15]. Several studies have demonstrated the involvement of MAGMAS in human diseases. A homozygous missense mutation in MAGMAS correlates with severe skeletal dysplasia [16]. MAGMAS mRNA shows increased expression in murine and rat pituitary adenomas cell lines [15, 17]. MAGMAS level is also upregulated in patient samples of prostate and ovarian cancers [18, 19]. We previously showed increased expression of MAGMAS in patient-derived glioblastoma tissues [20]. We further reported that a small molecule MAGMAS inhibitor, BT9, decreases cell proliferation, migration, and oxidative phosphorylation in human glioblastoma cell lines and can cross the blood brain barrier [20]. The aim of this study was to investigate whether the therapeutic effect of MAGMAS inhibition can be extended to pediatric medulloblastoma.

## Methods and Materials

### Cell Culture

The established SHH activated tp53 mutant, DAOY [21], and non-WNT/non-SHH (group3), D-425, pediatric medulloblastoma cell lines were cultured in DMEM/F-12 medium containing 10% FBS (Cat # 12306C, Sigma), and 1% Penicillin/streptomycin (Cat # 15140-122, Gibco). All cells were tested for mycoplasma infection and maintained at 37°C in a humidified incubator with 5% CO2. BT9 compound was synthesized as previously described [22] and dissolved in DMSO and Captisol (Ligand Pharmaceuticals, San Diego) for *in vitro* and *in vivo* studies, respectively.

### Cell viability assay

The cells were seeded at approximately 5× 10^3^/well in a final volume of 200 μl in 96-well microtiter plates and allowed to grow for 24 h before adding BT9 compound. 5mg/ml MTT solution (Cat # 50-213-524, Fisher Scientific) was added at 24h, 48, and 72h after BT9 treatment and cells were incubated at 37 °C for 4 hours. The culture medium was then aspirated and DMSO (200 μl/well, Fisher Scientific) was added to dissolve the dark blue formazan crystals. The absorbance was measured at a wavelength of 570 nm. Three independent experiments with 3 replicates per condition were performed. Data are expressed as relative survival compared to DMSO, and IC50 was determined using non-linear regression analysis on effect-log concentration curves.

### Apoptosis assay

DAOY apoptotic cells were detected by Click-iT™ TUNEL Assay in accordance with the manufacturer’s instructions (Invitrogen, Cat# C10247). 2.5 × 10^4^ cells were seeded in Nunc Lab-Tek chamber slides system (Fisher Scientific, Cat# 177445) and treated with daily BT9 for 24 and 48 hours. Slides were imaged with a Nikon Ti-E microscope and analyzed using Image J. More than 3000 nuclei were counted per field; the experiment was repeated three times.

### Wound closure/migration assay

Cells (6 × 10^5^) were seeded in a 6-well plate to form a confluent monolayer. A scratch wound was made in the center of the well by scraping it with P200 pipette tip. Cells were then washed with PBS, treated with BT9 compound, and incubated to allow cells to migrate into the space cleared by the scratch. Cell migration was assessed by capturing images from identical locations along each wound using EVOS X10 microscope at 0,4,8, and 12 hours after the scratch.

### Transwell Invasion assay

Invasion assays were performed using a 24-well Matrigel Invasion chamber (Corning, USA) containing an 8μm pore size polyethylene terephthalate (PET) membrane treated with Matrigel basement membrane matrix (Cat # 354480). DAOY and D425 (2.5 × 10^4^) cells were pretreated with 3uM BT9 and plated in the top chamber in serum-free medium. Cells were allowed to invade into the lower chamber (containing culture medium with 10% fetal bovine serum) for 22 hours. Then, the nonmigratory cells were removed from the upper side of the membrane, and the cells on the lower surface of the membrane were fixed and stained with 0.5% crystal violet. The number of migratory cells was determined by counting the cells that penetrated the membrane using Nikon Ti-E microscope. Each experiment was performed in triplicate (n = 3 in each experiment)

### Seahorse XF24 metabolic flux analysis

Seahorse XF Mito Stress Test Kit (Cat # 103015-100, Agilent, Santa Clara, USA) was used to measure Oxygen consumption rates (OCR, pmol/min) and extracellular acidification rates (ECAR, mpH/min). One day prior to the experiment, 6× 10^4^ cells/ well were seeded in a XF24 cell culture plate in complete DMEM/F12 media. For D425, the wells were first coated with poly lysine. On the day of the experiment, cells were washed and incubated in XF assay medium supplemented with 17.5mM glucose, 2.5mM glutamine, and 0.5mM Sodium pyruvate for 1 hour at 37 °C in 0% CO2. Mitochondrial inhibitors: oligomycin (1μM), FCCP (0.5 μM) and rotenone/actinomycin A (1μM) were added based on the manufacturer’s recommendation. After the experiment, all the cells were recovered, and OCR/ECAR measurements were normalized to protein content per well using BCA protein assay kit (Thermo Fisher Scientific).

### Intracranial Xenograft model

Xenograft of tumor cell suspension was carried out in 6–8-week-old NSG mice (NOD-SCID gamma mouse). Mice were obtained from the laboratory breeding colony, established by breeding pairs purchased from Jackson Laboratories. All procedures were reviewed and approved by UCI Institutional Animal Care Use Committee. Sample sizes were chosen to minimize the number of animals required to get significant results.

One week prior to the cell injection, animals were surgically implanted with a catheter into the jugular vein, as previously described [23]. Briefly, catheters were constructed with guide cannula bent at a curved right angle and fitted with 6cm length silastic tubing. For the intravenous surgery, the catheter port was subcutaneously implanted in the animal’s back, and tubing was guided under the skin at the shoulder/neck to the right jugular vein. Thereafter, 1cm of the tubing was inserted into the vein and secured with surgical silk suture [23]. On the day of stereotactic implantation, mice were anesthetized by isoflurane/oxygen vapor mixture. An incision of ∼1.5 cm was made along the mediolateral line near the back of the skull using a scalpel (Feather Safety Razor Co.). Once the area was clean, a burr hole was drilled using a handheld microdrill. D425 cell suspension (2μl; 1.75 ×10^5^ cells) were injected into the cerebellum of 19 mice using a 30G Hamilton Syringe (7803-07, Hamilton Company) with stereotaxic guidance (AP: - 2mm; DV: -2mm; ML: 1mm, relative to lambda). The needle was left in place for an additional minute to limit reflux along the injection. Following closure of the incision, mice were removed from isoflurane and injected intraperitoneally with 10 mg/kg carprofen and 10mg/kg enrofloxacin for 5 days. Seven days after the surgery, animals received 50 mg/kg BT9 three times per week until they were euthanized. Captisol was used as control.

### Statistical analysis

Statistical analysis was performed using GraphPad Prism 9 software. Group comparisons were performed using Student’s t-test or ANOVA based on whether two or more groups were compared, respectively. Results were expressed as means ± SEM. Mann-Whitney test was used for group comparisons (two-tailed). Survival analysis was performed using log-rank test. P < 0.05 was considered significant

## Results

### MAGMAS inhibitor reduces cell growth and induces cell death in both SHH-driven and group3 medulloblastoma cell lines

To test the effects of BT9 on cell growth and viability, we performed MTT assay on established medulloblastoma DAOY and D425 cell lines. The cells were treated with increasing concentration of BT9 at three different time points, 24, 48, and 72hrs. As shown in Figure 1A, BT9 treatment demonstrated a significant inhibition on DAOY cell growth in a dose- and time-dependent manner (24hrs IC50: 3.6uM, 48hrs IC50: 2.3uM, 72hrs IC50: 2.1uM). Similar sensitivities were observed on D425 cells (Fig.1B, 24hrs IC50: 3.4uM, 48hrs IC50: 2.2uM, 72hrs IC50: 2.1uM). To understand whether these reductions in cell growth occurred due to induction of cell death, we next performed TUNNEL assay. This analysis showed that BT9 led to a significant increase in the proportion of TUNNEL-positive cells (i.e., dying cells) in both DAOY (Fig2, A, B, and E; control 1.7% vs BT9 10.1%, p = 0.0004, 24hrs; 1.8% vs 35.3%, p <0.0001, 48hrs) and D425 (Fig2. C, D, F; control 1.4% vs BT9 15.3%, p = 0.0001, 24hrs; 19.8% vs 43.6%, p = 0.02, 48hrs) cells. Taken together, our data suggest that BT9 inhibits medulloblastoma cell growth by inducing cell death.

**Fig 1.**
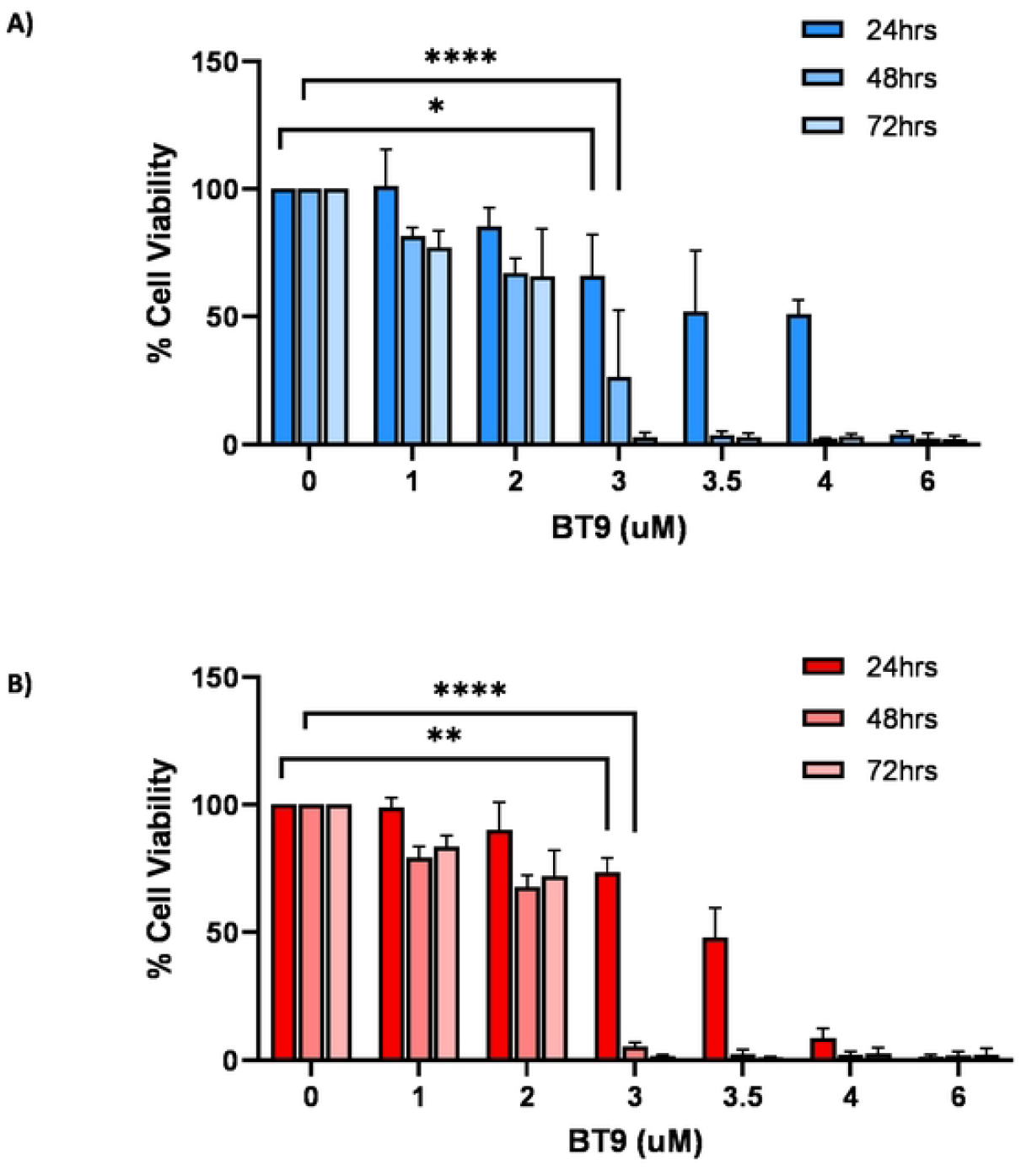
BT9 exhibits a dose- and time-dependent cytotoxic effect on medulloblastoma. DAOY **(A)** and D425 **(B)** were incubated for 24, 48, and 72 hours with increasing concentrations of BT9. Cell viability was measured by MTT assay. The relative numbers of proliferating cells compared with control are presented as the mean ± SEM. *p < 0.05, **p < 0.01, ****p < 0.0001.

**Fig 2.**
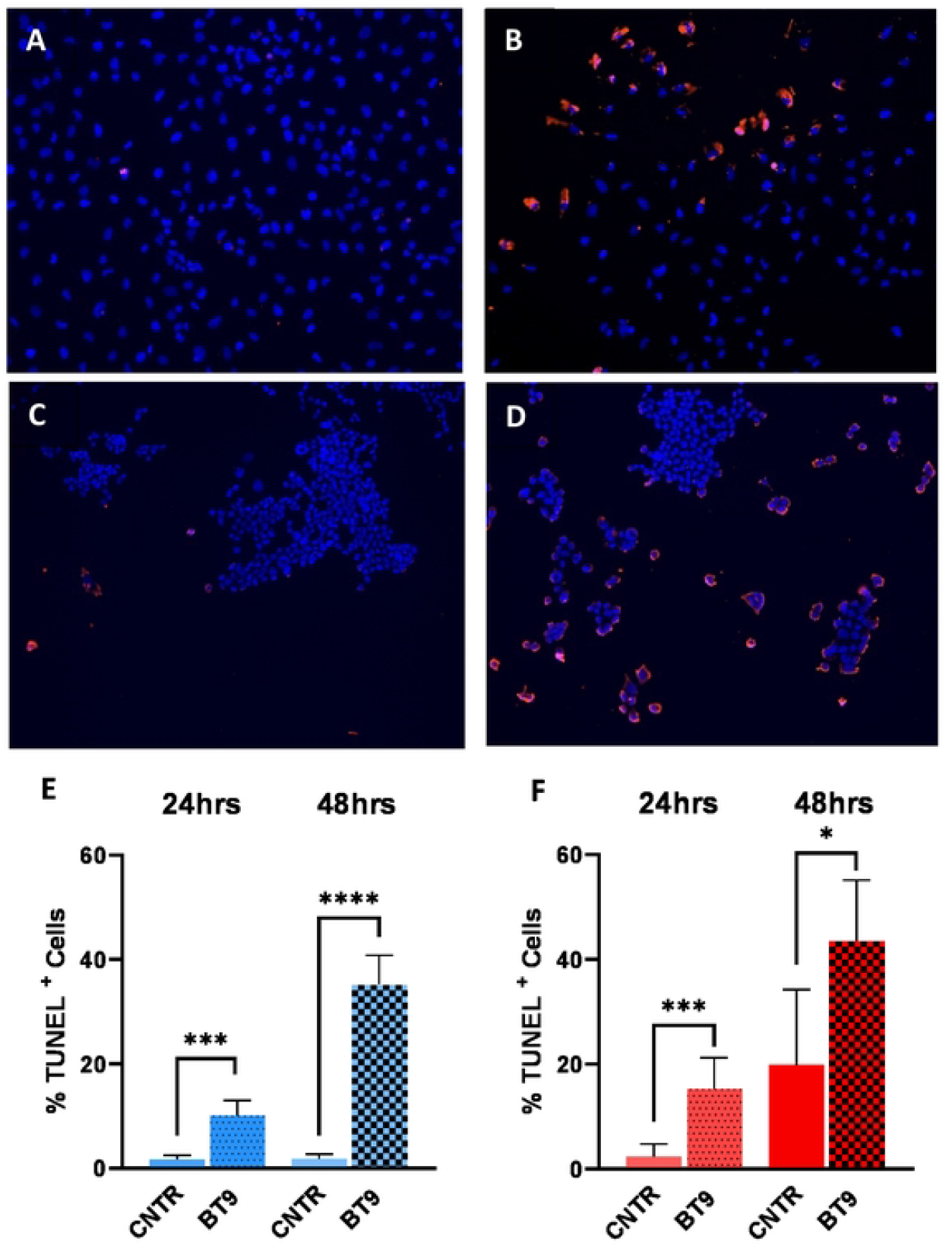
MAGMAS inhibition promotes medulloblastoma cell death. Apoptosis– based quantification of TUNEL-positive cells (dead cells) following BT9 treatment in DAOY (A, B, and E) and D425 (C, D, and F) cells after 24 and 48 hours (E and F). BT9 treatment significantly increases cell death in both cells (*p < 0.05, ***p < 0.001, ****p < 0.0001, BT9 vs vehicle control, n = 9). 10x magnification.

### BT9 inhibits medulloblastoma cell migration and invasion

We next investigated the effect of BT9 on medulloblastoma cell migration through scratch wound assays with DAOY cells. We were unable to perform this assay with D425 cells due to their nonadherent nature. Representative photographs (Fig. 3A-H) demonstrated control (Fig. 3A-D) versus 3uM-treated BT9 cells (Fig.3E-H) at 4, 8, and 12 hours after performing the scratch. Our results showed that BT9 reduced migration of DAOY cells to the center of the scratch wound. To better understand the effect of BT9 in inhibiting cell movement, we performed invasion assay with Matrigel-coated inserts, which mimics the natural barrier that cells must overcome to invade into other tissues.

**Fig 3.**
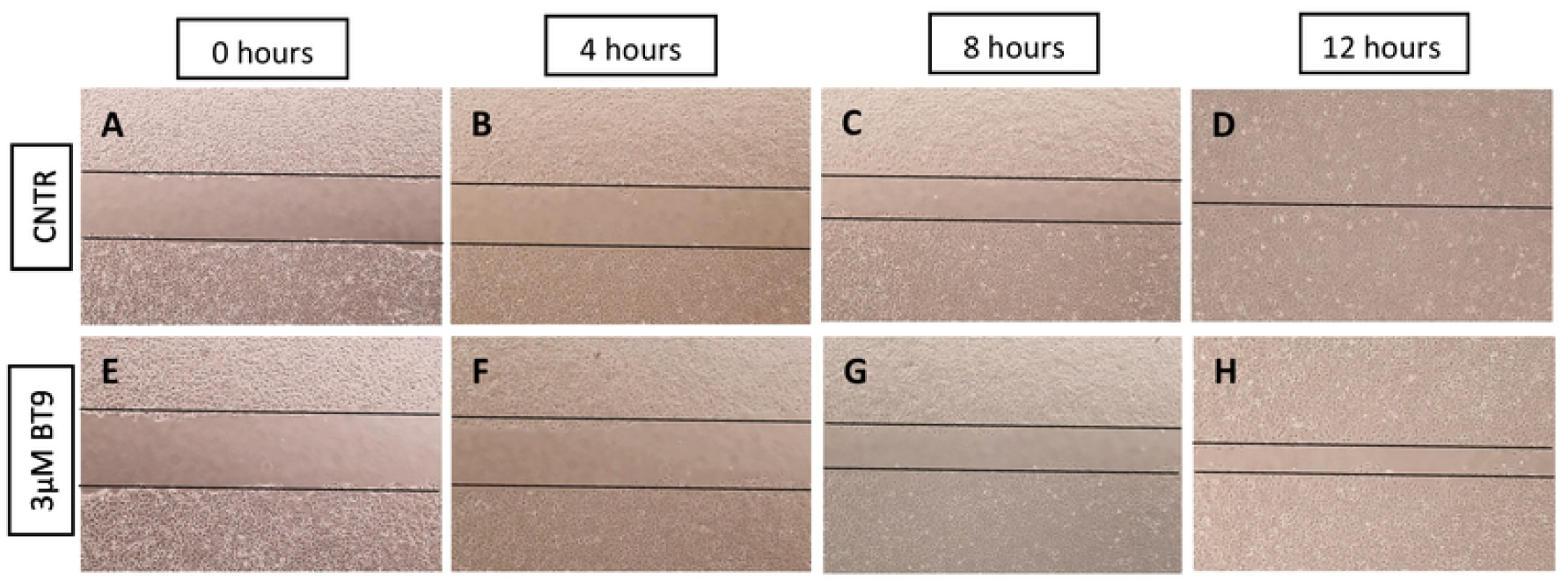
MAGMAS inhibition reduces DAOY cells migration. Migration of DAOY cells treated with vehicle control (A-D) or 3 µM BT9 (E-H) was assessed using the scratch wound assay. Representative photographs were taken immediately after the scratch (T0) and 4 hours intervals (T4, 8, and 12). 4X magnification.

Consistent with our migration data, we found that BT9 treatment led to a significant reduction in the invasive capacity of both DAOY and D425 cells after 22 hours (Fig. 4A-C). In total, our findings demonstrate that BT9 may provide an effective means of inhibiting medulloblastoma cell migration and invasion.

**Fig 4.**
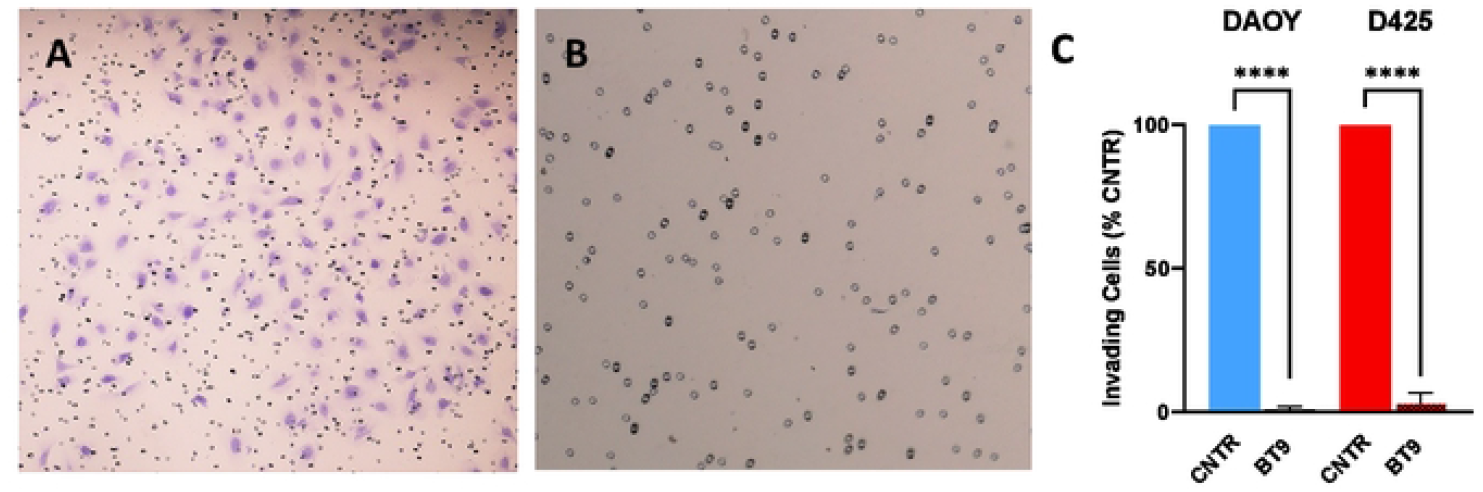
Invasion of medulloblastoma cells treated with vehicle or BT9 (3µM) was assessed using inserts coated with Matrigel matrix. (A and B) Representative images show the invading DAOY cells (purple cells). (C) BT9 treatment significantly decreased the number of both DAOY and D425 cells invading through the Matrigel matrix (****p < 0.0001, BT9 vs vehicle control, n = 6). 4x magnification.

### MAGMAS inhibition in medulloblastoma cell lines alters mitochondrial respiration function

It has been shown that MAGMAS promotes cellular tolerance toward oxidative stress by increasing electron transport chain (ETC) function that results in reduced ROS production [14]. To measure the effect of MAGMAS inhibition on ETC, we assessed basal cellular oxygen consumption rate (OCR) using the Seahorse Biosciences Extracellular Flux Analyzer. In D425 cells, BT9 treatment for 48hrs decreased basal respiration (p=0.006), ATP production (p=0.002), maximal respiration (p=0.005), and spare respiratory capacity (p=0.02) in a dose-dependent manner, suggesting that the oxidative phosphorylation (OXPHOS) activity has been reduced (Fig.5 A and B). We observed the same trend in DAOY cell line (data not shown).

**Fig 5.**
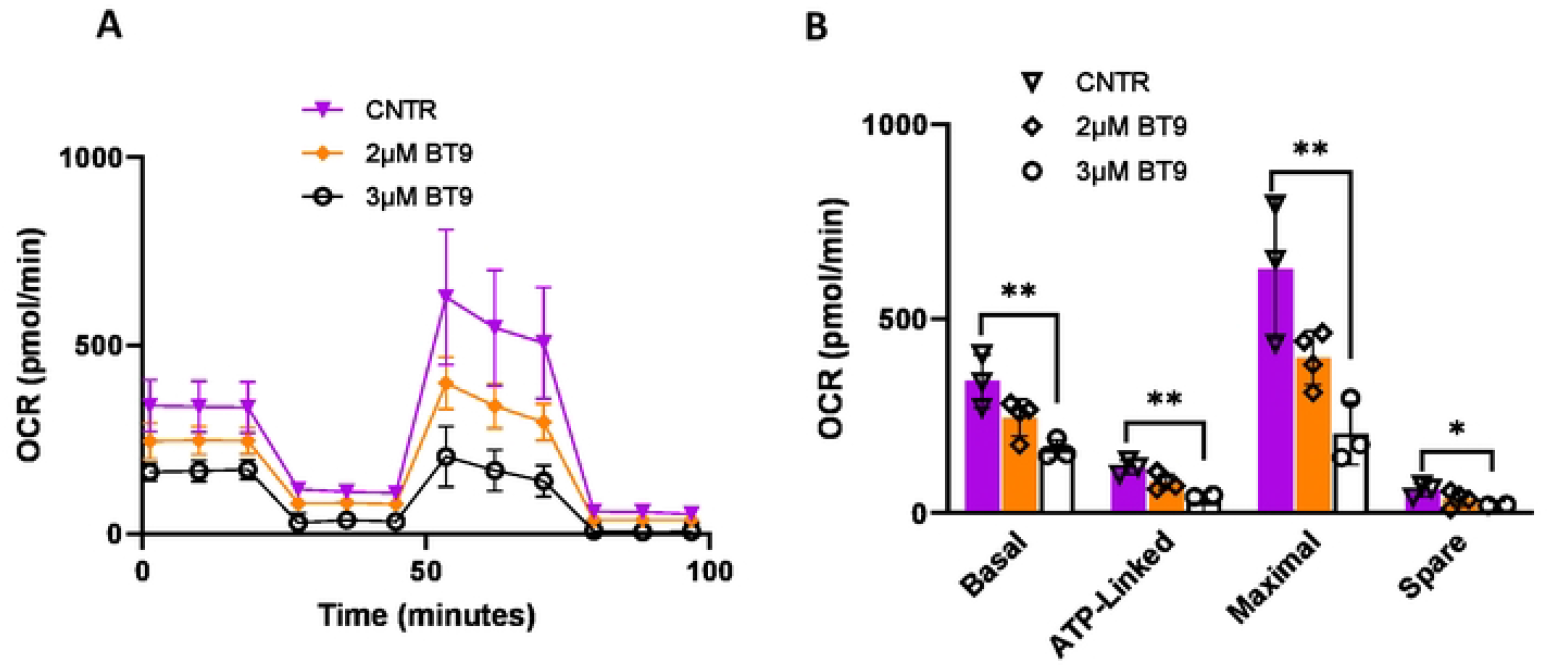
MAGMAS inhibition impaired mitochondrial respiration in D425 cells. Oxygen consumption rate (OCR) profile of D425 treated with 2μM and 3μM BT9 for 48 hours. (A) The result showed a significant reduction in oxygen consumption of D425 cells treated with 3µM BT9. (B) Bar chart showing the results of mitochondrial respiration changes in BT9-treated cells, which were analyzed with basal respiration, ATP production, maximal respiration, and spare respiratory capacity. (*p < 0.05, **p < 0.01, 3µM BT9 vs control.

### BT9 does not extend survival in a group3 medulloblastoma animal model

We investigated the efficacy of BT9 as monotherapy in the group3 medulloblastoma mouse model via intracranial injection of D425 cells. 7 days after cell implementation, 50mg/kg BT9 was given intravenously through the jugular vein, three times per week until the animals showed signs of cancer development and were euthanized (Fig. 6A). We did not observe any significant extension in median survival of the BT9 treated animals (Fig. 6B).

**Fig 6.**
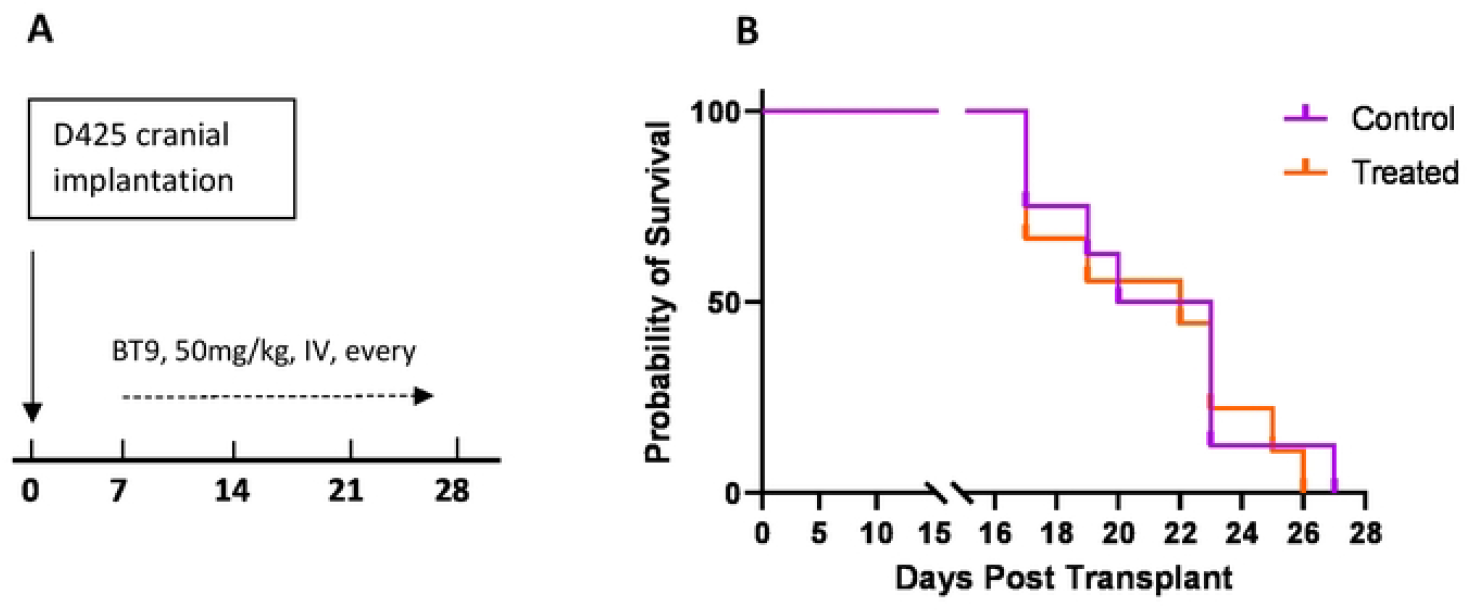
BT9 does not change survival in an orthotopic xenograft model of group3 medulloblastoma. A: Experimental setup. Immunodeficient (SCID) mice were intracranially implanted with D425 cells and treated every other day with an intravenous injection of 50 mg/kg BT9 or vehicle control (Captisol). BT9 treatment started 7 days after implantation. B: Kaplan-Meier survival analysis for *in vivo* experiment. BT9 treatment did not lead to a statistically significant increase in median survival compared to vehicle-treated controls (27 days; n = 8 control vs n= 9 treated group).

## Discussion

Mitochondria are essential organelles serving not only as energy factories but also as key players in numerous cell functions such as proliferation and death. Alterations in mitochondrial metabolic and signaling pathways have been associated with both tumorigenesis and cancer progression [24]. In recent years, mitochondria have emerged as a therapeutic target in many types of cancers, such as breast, lung, colon, and pancreas [24-26]. These agents target various mitochondrial functions such as metabolism, apoptotic pathways, and ROS homeostasis, but so far they have not shown any significant therapeutic effects [26]. BT9 is a novel small molecule that binds to MAGMAS and potentially blocks protein trafficking across the inner membrane, and it has shown inhibitory effects on proliferation and migration in glioblastoma and prostate cancer cell lines [20, 27]. In this study, we investigated BT9 efficacy in pediatric medulloblastoma.

We first showed that BT9 decreases cell growth in SHH-activated tp53 mutant and non-WNT/non-SHH (group3) medulloblastoma cells *in vitro*. Both DAOY and D425 cells showed the same sensitivity to MAGMAS inhibition (3.64 vs 3.38 µM at 24hrs, 2.28 vs 2.17 µM at 48hrs, and 2.14 vs 2.16 µM at 72hrs, respectively). These effects are consistent with our results in glioblastoma cells, where IC50 was found around 2.5 µM after 72 hrs BT9 treatment [20]. We also found an increased cell death in BT9-treated cells in a caspase-dependent manner, which is consistent with our previous results in glioblastoma cells [20]. However, Yang et al. reported both caspase-dependent and -independent pathways as the modes of cell death in the prostate cancer [27].

Measuring mitochondrial metabolic activity, our results showed that BT9 significantly decreases OXPHOS in D425 cells, suggesting increased production of ROS, which then results in oxidative stress and cell death. This is of clinical relevance since OXPHOS inhibitors have been shown to improve responses to anticancer therapy in certain cancers, such as melanomas, lymphomas, colon cancers, leukemias and pancreatic ductal adenocarcinoma [28]. We also showed that OCR baseline is lower in D425 (250pmol/min) compared to DAOY (620pmol/min) cells. These findings are consistent with recent data demonstrating that OXPHOS genes are the most downregulated ones in group3 medulloblastoma [29].

We further demonstrated that BT9 reduces medulloblastoma cell migration and invasion, which is crucial, given medulloblastoma’s unique propensity for metastasis [1, 30]. It has long been shown that ROS may activate different processes associated with migration and invasion through cytoskeleton remodeling [31, 32]. ROS may promote tumor cell invasion by stimulation of the proteolytic degradation of extracellular matrix components such as glycosaminoglycan leading to metastasis [32]. Furthermore, increased ROS stimulates epithelial-mesenchymal transition through the activation of NF-κB and TGF-β signaling [32].

Despite promising *in vitro* results, we did not find any significant increase in survival in D425 medulloblastoma *in vivo* model after BT9 treatment. This result does not completely negate the efficacy of BT9 in the *in vivo* system. It simply shows that under the protocol used in this study, BT9 is not effective, and further work needs to be optimized. We suggest using higher (>50mg/kg) and/or more frequent (ex. daily) doses. We started BT9 treatment 7 days after cell implantation. One might try to administer the drug at earlier time points. It is possible that the tumor has already progressed into a late-stage cancer by the first date of our treatment. Since BT9 will probably be used as one component of a broader treatment regimens, one might plan to investigate BT9 in combination therapy with current treatment modalities. In fact, we found that BT9 shows additive effects when combined with temozolomide in glioblastoma cells (unpublished data). Finally, this work ideally should be validated in patient-derived xenograft models before moving to the clinic.

## Acknowledgement

DAOY and D425 cells were generous gifts from Dr. Wechsler-Reya (Columbia University) and Dr. Cheshier (University of Utah) laboratories, respectively. The authors thank Dr. Kenney’s lab and Marilyn Chwa at UCI for the training and providing the reagents for the Seahorse assay. This work is supported by NINDS R01 (NS109423) to Drs. Daniela Bota and Bhaskar Das, 3R01 NS109423-02 Supplement to Javier J. lepe, UCI Cancer Center Award number P30CA062203 from the National Cancer Institute, Pediatric Cancer Research Foundation, Chao Family Comprehensive Cancer Center, and CHOC Neuroscience Institute. We also thank Coon family, Kevin Freeman family foundation, and Luhmann Family Trust donation for generous gifts to this project.

## References

1. Juraschka, K. and M.D. Taylor, Medulloblastoma in the age of molecular subgroups: a review. J Neurosurg Pediatr, 2019. 24(4): p. 353–363.

2. Archer, T.C., E.L. Mahoney, and S.L. Pomeroy, Medulloblastoma: Molecular Classification-Based Personal Therapeutics. Neurotherapeutics, 2017. 14(2): p. 265–273.

3. Martin, A.M., et al., Management of pediatric and adult patients with medulloblastoma. Curr Treat Options Oncol, 2014. 15(4): p. 581–94.

4. Orr, B.A., Pathology, diagnostics, and classification of medulloblastoma. Brain Pathol, 2020. 30(3): p. 664–678.

5. Roussel, M.F. and G.W. Robinson, Role of MYC in Medulloblastoma. Cold Spring Harb Perspect Med, 2013. 3(11).

6. Kim, J.W., et al., Evaluation of myc E-box phylogenetic footprints in glycolytic genes by chromatin immunoprecipitation assays. Mol Cell Biol, 2004. 24(13): p. 5923–36.

7. Li, F., et al., Myc stimulates nuclearly encoded mitochondrial genes and mitochondrial biogenesis. Mol Cell Biol, 2005. 25(14): p. 6225–34.

8. Popay, T.M., et al., MYC regulates ribosome biogenesis and mitochondrial gene expression programs through its interaction with host cell factor-1. Elife, 2021. 10.

9. Sinha, D., et al., Role of Magmas in protein transport and human mitochondria biogenesis. Hum Mol Genet, 2010. 19(7): p. 1248–62.

10. Morrish, F. and D. Hockenbery, MYC and mitochondrial biogenesis. Cold Spring Harb Perspect Med, 2014. 4(5).

11. Whitfield, J.R. and L. Soucek, The long journey to bring a Myc inhibitor to the clinic. J Cell Biol, 2021. 220(8).

12. Gebert, N., et al., Mitochondrial protein import machineries and lipids: a functional connection. Biochim Biophys Acta, 2011. 1808(3): p. 1002–11.

13. Mokranjac, D., et al., Structure and function of Tim14 and Tim16, the J and J-like components of the mitochondrial protein import motor. EMBO J, 2006. 25(19): p. 4675–85.

14. Srivastava, S., et al., Magmas functions as a ROS regulator and provides cytoprotection against oxidative stress-mediated damages. Cell Death Dis, 2014. 5(8): p. e1394.

15. Tagliati, F., et al., Magmas overexpression inhibits staurosporine induced apoptosis in rat pituitary adenoma cell lines. PLoS One, 2013. 8(9): p. e75194.

16. Mehawej, C., et al., The impairment of MAGMAS function in human is responsible for a severe skeletal dysplasia. PLoS Genet, 2014. 10(5): p. e1004311.

17. Tagliati, F., et al., Magmas, a gene newly identified as overexpressed in human and mouse ACTH-secreting pituitary adenomas, protects pituitary cells from apoptotic stimuli. Endocrinology, 2010. 151(10): p. 4635–42.

18. Ahmed, N., et al., Ovarian Cancer, Cancer Stem Cells and Current Treatment Strategies: A Potential Role of Magmas in the Current Treatment Methods. Cells, 2020. 9(3).

19. Jubinsky, P.T., et al., Magmas expression in neoplastic human prostate. J Mol Histol, 2005. 36(1-2): p. 69–75.

20. Di, K., et al., Magmas inhibition as a potential treatment strategy in malignant glioma. J Neurooncol, 2019. 141(2): p. 267–276.

21. Naeem, A., et al., Regulation of Chemosensitivity in Human Medulloblastoma Cells by p53 and the PI3 Kinase Signaling Pathway. Mol Cancer Res, 2022. 20(1): p. 114–126.

22. Jubinsky, P.T., et al., Design, synthesis, and biological activity of novel Magmas inhibitors. Bioorg Med Chem Lett, 2011. 21(11): p. 3479–82.

23. Fowler, C.D. and P.J. Kenny, Intravenous nicotine self-administration and cue-induced reinstatement in mice: effects of nicotine dose, rate of drug infusion and prior instrumental training. Neuropharmacology, 2011. 61(4): p. 687–98.

24. Missiroli, S., et al., Cancer metabolism and mitochondria: Finding novel mechanisms to fight tumours. EBioMedicine, 2020. 59: p. 102943.

25. Ghosh, P., et al., Mitochondria Targeting as an Effective Strategy for Cancer Therapy. International Journal of Molecular Sciences, 2020. 21(9).

26. Neuzil, J., et al., Classification of mitocans, anti-cancer drugs acting on mitochondria. Mitochondrion, 2013. 13(3): p. 199–208.

27. Yang, J., et al., Magmas Inhibition in Prostate Cancer: A Novel Target for Treatment-Resistant Disease. Cancers (Basel), 2022. 14(11).

28. Zhao, Z., et al., The Effect of Oxidative Phosphorylation on Cancer Drug Resistance. Cancers (Basel), 2022. 15(1).

29. Li, Q., et al., Mitochondrial subtype MB-G3 contains potential novel biomarkers and therapeutic targets associated with prognosis of medulloblastoma. Biomarkers, 2023. 28(7): p. 643–651.

30. Northcott, P.A., et al., Medulloblastoma. Nat Rev Dis Primers, 2019. 5(1): p. 11.

31. Yanes, B. and E. Rainero, The Interplay between Cell-Extracellular Matrix Interaction and Mitochondria Dynamics in Cancer. Cancers (Basel), 2022. 14(6).

32. Denisenko, T.V., A.S. Gorbunova, and B. Zhivotovsky, Mitochondrial Involvement in Migration, Invasion and Metastasis. Front Cell Dev Biol, 2019. 7: p. 355.

